# The chromatin conformation landscape of Alzheimer’s disease

**DOI:** 10.1101/2024.04.09.588722

**Authors:** R Nativio, Y Lan, G Donahue, O Shcherbakova, N Barnett, KR Titus, H Chandrashekar, JE Phillips-Cremins, NM Bonini, SL Berger

**Affiliations:** Department of Brain Sciences and the UK Dementia Research Institute, Imperial College London, London, UK; Epigenetics Institute, Perelman School of Medicine, University of Pennsylvania; Philadelphia, PA; Department of Cell and Developmental Biology, Perelman School of Medicine, University of Pennsylvania; Philadelphia, PA; Department of Biology, School of Arts and Sciences, University of Pennsylvania; Philadelphia, PA; Department of Genetics, Perelman School of Medicine, University of Pennsylvania; Philadelphia, PA; Department of Bioengineering, School of Engineering and Applied Science, University of Pennsylvania, Philadelphia, PA

**Author notes:** Co-corresponding authors: R. Nativio, NM Bonini, SL Berger.

## Abstract

We have been investigating epigenetic alterations in the brain during human aging and Alzheimer’s disease (AD), and have evidence for histone acetylation both protecting the aging epigenome and driving AD. Here we extend our studies to chromatin architecture via looping studies, and with binding studies of key proteins required for looping: CTCF and RAD21. We detected changes in CTCF and RAD21 levels and localization, finding major changes in CTCF in AD compared to fewer changes in healthy aging. In our study of 3D genome conformation changes, we identified stable topological associating domains (TADs) in Old and AD; in contrast, in AD, there is loss of interaction at genomic sites/loops within TADs, likely reflecting the loss of CTCF. We identified genes and potential transcription factor binding at the loops that are lost in AD. in addition, we found enrichment of CTCF peak losses for AD eQTLs, suggesting that architectural dysfunction has a role in Alzheimer’s. Functional experiments lowering the homologues of several key genes in a *Drosophila* model of Aβ42 toxicity exacerbate neurodegeneration. Taken together, these data indicate both functional protections and losses occur in the Alzheimer’s brain genome compared to normal aging.

## INTRODUCTION

Epigenetics is the study of changes in the genome beyond changes in the DNA sequence; for example, methylation changes and histone modifications ^1–4^. The genome is also not thought to be just a linear sequence of DNA and associated proteins, but to be highly organized into domains and structural regions that are critical to proper gene expression ^5–7^. We are only beginning to understand how these higher organizational elements of the genome impact normal gene expression and maintenance of proper cellular state. These structural features of the genome are fascinating, yet understanding of their functional impact if disrupted or altered with disease, is only now becoming addressed.

Of brain disease, Alzheimer’s disease is among those requiring most urgent attention, given the increased incidence with age. Alzheimer’s disease leads to a deterioration of brain cognitive function, and ultimately loss of brain cells. To understand the changes that may occur to the genome with disease, our work ^8,9^ and others ^10–13^ has focused on epigenetic status of the brain in Alzheimer’s. Alzheimer’s is initially primarily characterized by a loss of memory, with a focus on the temporal and hippocampal regions of the brain.

CCCTC-binding factor (CTCF) and cohesin are architectural proteins which partition the genome in fundamental chromatin conformation structures such as Topologically Associating Domains (TADs) ^5,14,15^. Inside TADs, CTCF and cohesin also mediate long-range chromatin interactions, including linking enhancers to promoters for proper gene expression. CTCF also functions as a transcription factor to guide gene expression and insulator so that the proper gene, and not neighboring genes, are activated. The insulation function may be attributed to its role in chromatin conformation, potentially by establishing silencing domains, as well as through other mechanisms ^16^. Additional chromatin factors, such as Mediator and YY1, also facilitate long-range chromatin interactions.

We previously identified histone acetyl changes at non-coding regions of the genome in AD and with age, considered to be in enhancers ^8,9^. However, changes in the 3D topology of the genome, and particularly in enhancer-promoter interactions, could be involved in the mechanisms underlying gene dysfunction in AD. To address whether higher order changes occur in AD, we investigated chromatin factors CTCF and RAD21, and their related loops with normal aging and compared to AD. We considered that integrating this higher order understanding of the genome may reveal greater insight into how the genome deteriorates in AD, and therefore what types of approaches could be used to stabilize the genome for therapeutic advancement. Here we used CTCF and RAD21 ChIP-seq, HiC and CTCF HiChIP to probe the features of the AD genome relative to young and old, using tissue from the memory center. We also performed H3K9me3 and H3K27me3 to show that silencing also shows some changes between control and AD. Probing a *Drosophila* in vivo model highlighted that some of these interactions are critical to protect the brain from degeneration. These studies provide greater insight into the functional changes to the genome in AD that may contribute to disease.

## RESULTS

### Differential enrichment of CTCF and cohesin at unique genomic loci in Alzheimer’s disease compared to healthy aging

To investigate the role that chromatin conformation and associated proteins may play in Alzheimer’s disease (AD), we performed CTCF and RAD21 ChIP-seq, HiC and CTCF HiChIP across a set of Alzheimer’s affected (defined as AD; N=11), healthy age-matched (defined as Old; N=10) and healthy younger (defined as Young; N=9) controls from patient samples that we previously investigated for H3K27ac, H3K9ac, H3K122ac and H4K16ac changes genome-wide ^8^. Healthy younger samples were included to investigate age-related changes, to distinguish from disease-related changes. Tissue samples were derived from the lateral temporal lobe, a region affected relatively early in AD etiology, but retaining greater molecular integrity than other affected regions such as the entorhinal cortex.

CTCF and RAD21 ChIP-seq data were initially analyzed in the linear space, followed by investigation in the 3D space through integration with HiC and CTCF HiChIP data. ChIP-seq libraries were processed as in Nativio et al. ^8,9^, with peaks detected using MACS2 in pooled samples from each study group (**Fig 1A**). Subsequently, differential peak enrichment analysis was conducted using reads from individual samples. To address potential confounding variables, such as changes in cell type proportions, the top 10% of peaks with the highest Pearson’s correlation with sample neuronal fraction were masked. Neuronal fractions were previously quantified for each sample by flow cytometry of NeuN-stained nuclei^8,9^.

**Fig. 1:**
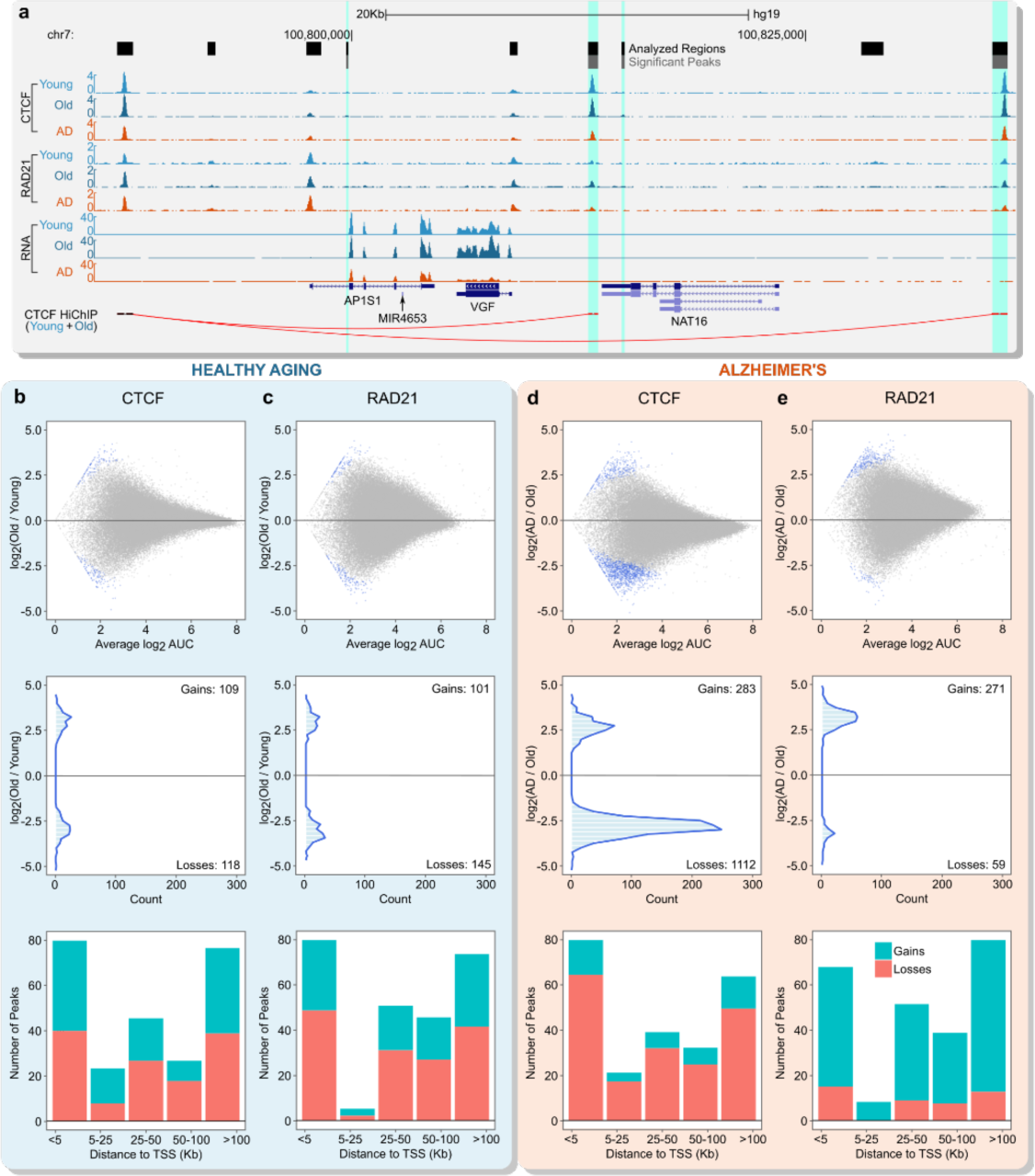
CTCF and RAD21 are differentially enriched in Alzheimer’s disease (AD) compared to healthy aging. **(a)** UCSC Genome Browser representation of CTCF, RAD21, CTCF loops and RNA-seq at a genomic region on chromosome 7 in Young, Old and AD. CTCF loops with reduced CTCF (highlighted in light blue) are shown to partition *AP1S1* and *VGF* into a looped domain. *AP1S1* and *VGF* expression is reduced in AD vs Old (q < 0.05). **(b)** Representation of CTCF changes in healthy aging (Old vs Young) showing: (*top*) scatterplot of peak fold-change vs average AUC (area under the curve) with blue dots representing peaks with significant fold-change (q < 0.05); (*center*) histogram showing the number of peaks with significant changes (q < 0.05) based on the fold-change enrichment; (*bottom*) Stacked barplot showing the number of CTCF peaks with significant changes (gains or losses) based on the distance from the TSS (Kb). **(c)** Representation of RAD21 changes in healthy aging (Old vs Young) showing: (*top*) scatterplot of peak fold-change vs average AUC (area under the curve) with blue dots representing peaks with significant fold-change (q < 0.05); (*center*) histogram showing the number of peaks with significant changes (q < 0.05) based on the fold-change enrichment; (*bottom*) Stacked barplot showing the number of RAD21 peaks with significant changes (gains or losses) based on the distance from the TSS (Kb). **(d)** Representation of CTCF changes in Alzheimer’s (AD vs Old) showing: (*top*) scatterplot of peak fold-change vs average AUC with blue dots representing peaks with significant fold-change (q < 0.05); (*center*) histogram showing the number of peaks with significant changes (q < 0.05) based on the fold-change enrichment; (*bottom*) Stacked barplot showing the number of CTCF peaks with significant changes (gains or losses) based on the distance from the TSS (Kb). **(e)** Representation of RAD21 changes in Alzheimer’s (AD vs Old) showing: (*top*) scatterplot of peak fold-change vs average AUC with blue dots representing peaks with significant fold-change (q < 0.05); (*center*) histogram showing the number of peaks with significant changes (q < 0.05) based on the fold-change enrichment; (*bottom*) Stacked barplot showing the number of RAD21 peaks with significant changes (gains or losses) based on the distance from the TSS (Kb).

To obtain a comprehensive understanding of how CTCF and RAD21 are related in healthy aging and AD, we first investigated their co-localization on the genome. Peak overlap analysis, considering all peaks detected in the three study groups (union of Young, Old and AD peaks), revealed that 76% of CTCF peaks overlapped with RAD21 (**Fig. S1A**). This overlap increased to 87% when focusing on peaks near transcription start sites (TSS) (within 1 Kb) (**Fig. S1A**). Conversely, RAD21 peaks showed fewer overlaps with CTCF, at 60%, decreasing further to 55% when examining regions located more than 1 Kb away from TSS, including enhancers (**Fig. S1A**). This analysis indicates that while CTCF and RAD21 overlap genome-wide, RAD21 is also enriched at genomic sites independently of CTCF, consistently with findings from other cell types ^17^.

Next, we performed differential peak enrichment analysis investigating changes in both healthy aging and AD. Differential peak-enrichment analysis in healthy aging (Old vs Young) indicated overall stable CTCF and RAD21 binding during normal aging (**Fig 1B-C**): there were only 109 peaks with CTCF gains and 118 peaks with CTCF losses (q < 0.05) (**Fig. 1B**) in Old vs Young; and 101 peaks with RAD21 gains and 145 with RAD21 losses (q < 0.05) (**Fig. 1C**). These changes were localized throughout the genome although with a preference for near the TSS (< 5 Kb from TSS) compared to other genomic regions (**Fig. 1B-C bottom**). In contrast, the analysis of CTCF and RAD21 changes in Alzheimer’s (AD vs Old) revealed a much higher number of changes (**Fig 1D-E**). There were 1,395 CTCF peak changes (q < 0.05) in AD vs Old, of which 1,112 were depleted and 283 were enriched in CTCF (q < 0.05) (**Fig. 1D**).

The analysis of RAD21 peaks revealed a starkly different trend of changes in AD (**Fig 1E**). First, there were overall fewer changes in RAD21 (n= 330) (q < 0.05) and second, 271 were enriched and only 59 were depleted in AD vs Old (q < 0.05) (**Fig. 1E**), revealing an opposite trend to CTCF changes. However, both CTCF and RAD21 changes were more prominent close to the TSS (**Fig. 1D-E bottom**). Given the differential trends in CTCF and RAD21 changes in AD, we investigated the level of overlap between these changes. Unexpectedly, there was almost no peak overlap between CTCF and RAD21 changes (**Fig. S1B**). Further quantitative comparison of CTCF and RAD21 enrichments at sites of differential peak detection corroborated these findings (**Fig. S1C-D**). Interestingly, sites with CTCF or RAD21 changes did not have significant changes in histone acetyl marks, indicating stable levels of open chromatin (**Fig. S1C-D**). These results indicate an overall stability of CTCF and RAD21 during normal aging, with prominent CTCF losses and modest RAD21 gains in AD. The distinct pattern of RAD21 changes observed in AD, despite the limited changes in peak numbers, implies that RAD21 may have a distinct role from CTCF, at least in AD.

Given the inclusion of the Young samples, we also investigated CTCF and RAD21 levels in Young at genomic sites displaying CTCF or RAD21 changes in AD vs Old (**Fig. S1C-D**). Our analyses indicated that the observed CTCF losses in AD vs Old were largely stable during healthy aging (Old vs Young) (**Fig. S1C**). In contrast, the identified RAD21 gains in AD showed trends opposite to those observed in healthy aging (**Fig. S1D**). We extended these analyses to the CTCF gains or RAD21 losses in AD vs Old, although few peak changes were observed for these classes of changes (trends reported in **Fig. S1E-F**). Most importantly, these findings emphasize that the CTCF losses observed in AD, which constitute the primary trends in these analyses, are distinctly linked to the disease.

Given that CTCF and RAD21 could influence gene expression indirectly by mediating enhancer-promoter interactions, and that CTCF also functions as a transcription factor, we investigated whether the enrichments of CTCF and RAD21 peaks correlated with gene expression at the genome-wide level (**Fig. S2A-H**). RNA-seq data from these samples were generated in our previous study ^8^. Surprisingly, there was no correlation between the levels of CTCF or RAD21 and the expression of the nearest gene in either Old or AD (**Fig. S2A-B and Fig. S2E-F**), even when analyzing significantly differentially expressed genes (q < 0.05) (**Fig. S2C-D and Fig. S2G-H**). While it is recognized that histone acetylation correlates with transcription, it is unclear whether the levels of a specific transcription factor directly correlate with gene expression. Therefore, we investigated whether CTCF and RAD21 indirectly impact the correlation between histone acetylation levels, particularly H3K27ac and gene expression. Our analysis focused on H3K27ac-enriched promoter (≤ 5 Kb from TSS) and enhancer (> 5 Kb from TSS) regions separately. The presence of CTCF or RAD21 at these specific sites enhanced the genome-wide correlations of H3K27ac with the expression of the nearest gene in AD (**Fig. S2I-P**). These analyses suggest that the involvement of CTCF and RAD21 in AD may be attributed to their indirect influence on gene expression, particularly through their mediation of enhancer-promoter interactions.

### Stable TADs are detected in healthy aging and AD

The changes we observed in CTCF and RAD21 in AD could involve changes in three-dimensional (3D) genome conformation. To examine this, we performed HiC and CTCF HiChIP in a subset of samples. HiC was performed to investigate 3D genome structures at the higher order level such as TADs, while CTCF HiChIP was performed to confirm the main findings from HiC and to analyze CTCF-mediated chromatin interactions. HiC and CTCF HiChIP data were then integrated to evaluate chromatin interaction changes at sites with CTCF losses.

HiC was performed on a subset of samples from Young, Old and AD groups, and the data processed following the method described in Zhang et al. ^18^ (**Fig. 2A**). We detected TADs in each study group using previous methods ^19,20^, and performed subsequent genome-wide characterization analyses (**Fig. S3A-C**). We examined the size distribution of TADs (**Fig. S3A**), the number of genes within TADs (**Fig. S3B**), and the number of TADs containing promoters, genes and other genomic elements across Young, Old and AD (Fig. S3C). The results revealed similar genomic features across all three study groups, which were also reflected at the chromosome level (**Fig. 2B-D**). Considering the overall similarity in 3D contact maps across the study groups (**Fig. 2B-D**), indicating no new formation or loss of TADs in AD or during healthy aging, we investigated potential differences in the intensity of TADs. For this analysis, we correlated the levels of HiC signal within TADs between AD and Old (**Fig. 2F**), as well as between Old and Young (**Fig. S4A**). This analysis demonstrated a significant level of correlation (Pearson r = 1 and p < 2.2e-16 for both comparisons), indicating minimal changes in TAD intensity. Overall, these analyses suggest that TADs remain stable throughout the process of healthy aging and in AD.

**Fig. 2:**
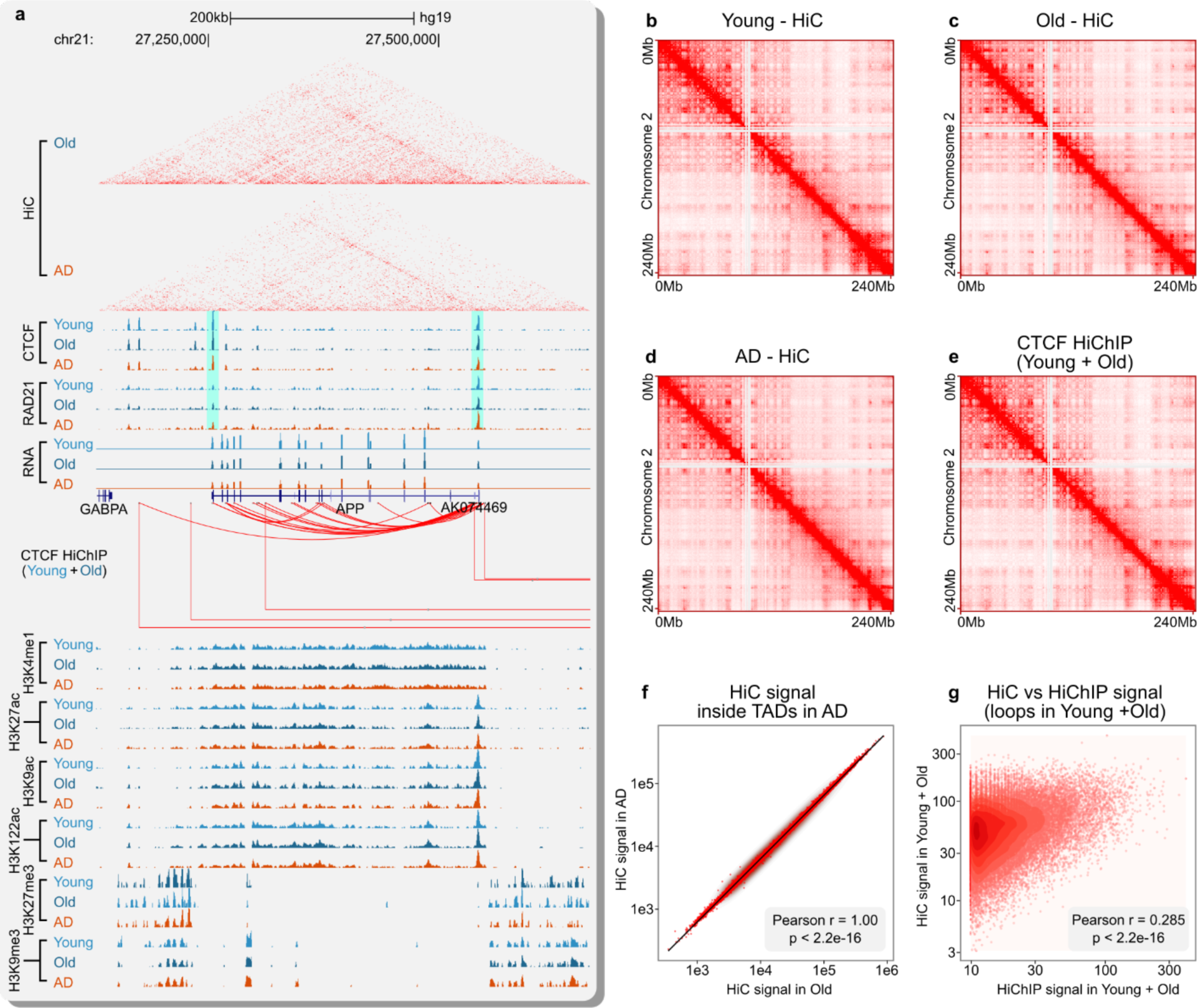
TADs are stable in healthy aging and AD. **(a)** UCSC Genome Browser representation of HiC, CTCF, RAD21, RNA and CTCF loops at the APP locus (chr21) in Young, Old and AD. Reported are also ChIP-seq tracks for H3K4me1, H3K27ac, H3K9ac, H3K122ac, H3K27me3 and H3K9me3 ChIP-seq generated from the same samples. Notably, the APP locus is situated within loops with CTCF binding at both anchor sites (highlighted in light blue). **(b-d)** HiC heatmap at chromosome 2 in **(b)** Young, **(c)** Old, **(d)** AD. **(e)** CTCF HiChIP heatmap at chromosome 2 (merged reads from Young and Old). **(f)** Scatterplot showing HiC signal in AD vs Old considering TADs detected in AD. **(g)** Scatterplot showing HiC vs CTCF HiChIP signal considering loops detected in HiChIP. Pearson’s correlation coefficient and p-value are indicated for panels f and g.

Next, we delved into the investigation of 3D structures dependent on CTCF. CTCF HiChIP was performed on samples from Young and Old (N = 2 per group) and the sequencing reads for the two groups were pooled. This pooling was carried out as no changes were observed in CTCF during aging, and it aimed to enhance the resolution of the subsequent analysis due to the increased sequencing depth. The HiChIP data were processed using reported methodologies ^21–23^. Loops were detected using both HICCUPS and FitHiChIP, leveraging their unique parameters. Following this, only the loops identified by both methods were retained for downstream analyses to ensure robustness and accuracy.

Three-dimensional interaction maps were visualized on a per chromosome basis (**Fig. 2E**), similar to the HiC data (**Fig. 2B-D**). We observed a high level of similarity between CTCF HiChIP (**Fig. 2E**) and HiC (**Fig. 2B-D**) upon visually comparing the 3D contacts at the chromosome level. There was also a significant correlation between loop intensity and HiC signal at corresponding sites (**Fig. 2G**). This finding aligns with the recognized role of CTCF in shaping the genome topology, including TADs, and it suggests that CTCF-mediated 3D topologies at the Mb level remain stable in AD, thereby corroborating the results from the HiC analysis.

### Long-range chromatin interactions are reduced at sites with CTCF losses

Following the identification of stable TADs in AD, we investigated whether changes in 3D genome conformation occurred at a smaller scale, particularly regarding enhancer-promoter interactions. To do so, we initially superimposed the identified CTCF loops onto TADs and categorized them based on their relative positions. Four main types of loops were identified (**Fig. 3A**): TAD loops (n=42), which encompass TADs; Boundary loops (n=1,991), where one CTCF anchor is located at the TAD boundary and the other within the TAD; Intra-TAD loops (n=27,607), with both anchors located within a TAD; and Non-TAD loops (n=1,378), where both anchors are outside a TAD. Genomic analysis of these loops (**Fig. S3D-F**) showed that, apart from the TAD loops, which were expectedly larger in size and had higher intensity, the Boundary, Intra-TAD, and Non-TAD loops exhibited similar size distribution and loop intensity (**Fig. S3D-E**). Notably, the Intra-TAD loops, constituting the majority of the loops, harbored the highest number of genes (**Fig. S3F**), suggesting their likely involvement in mediating enhancer-promoter interactions. The same class of loops also had the highest number of CTCF peak changes identified in AD at the corresponding anchor sites (**Fig. 3A**).

**Fig. 3:**
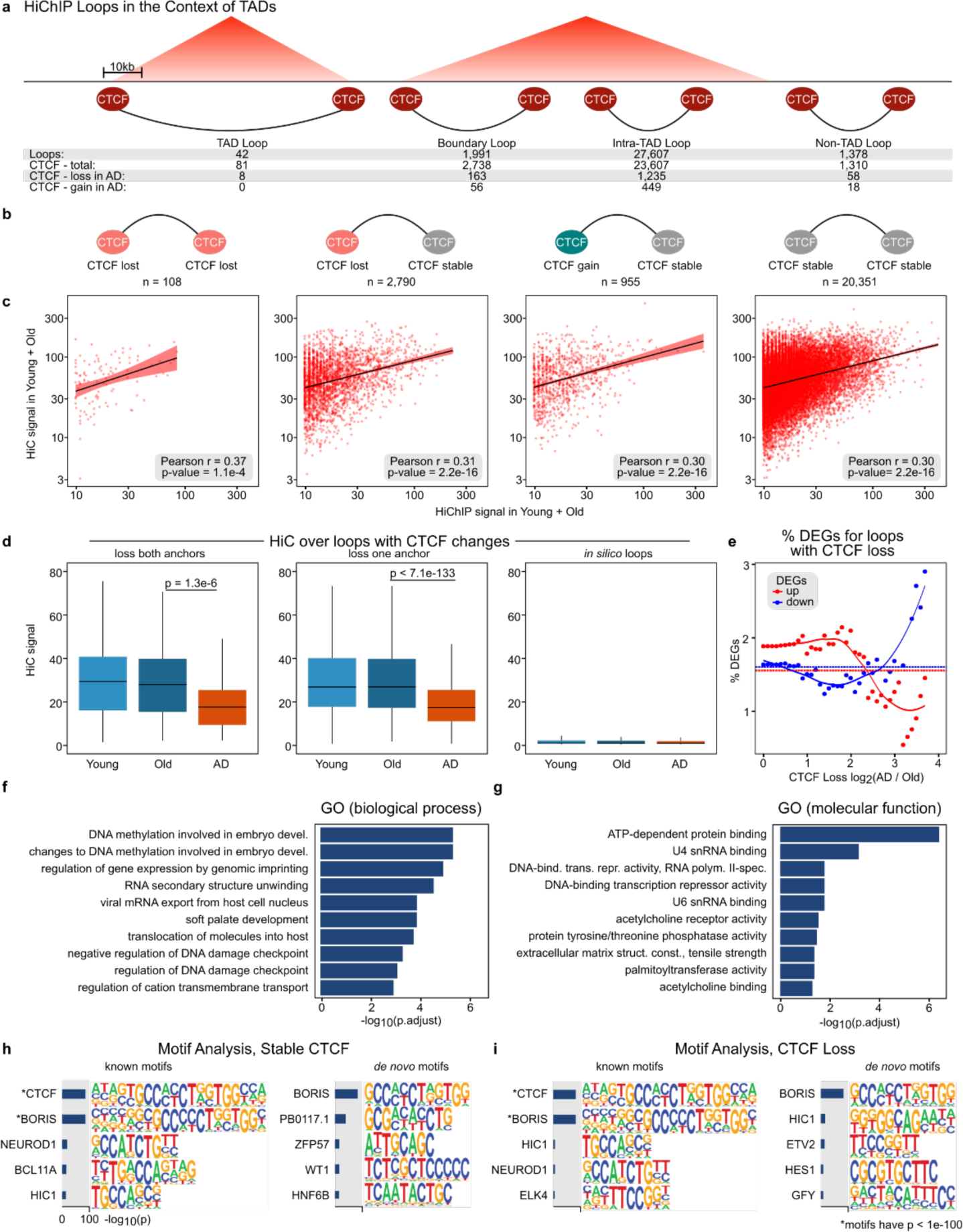
Reduction in 3D interactions at loops with CTCF losses in AD. **(a)** Graphical representation of CTCF loops based on their location relative to TADs and TAD boundaries. The number of CTCF peaks (total, loss or gained in AD vs Old) and loops is reported for each loop category. **(b)** Categorization of CTCF loops based on CTCF anchor changes in AD (CTCF loss at both ends, CTCF loss at one end, CTCF gained at one end, loops with stable CTCF). The number of loops for each category is represented underneath. **(c)** Scatterplot showing HiC vs CTCF HiChIP signal considering loops detected in HiChIP (Young + Old merged reads). Pearson’s correlation coefficient and p-value are indicated for each panel. **(d)** Boxplot showing HiC signal in Young, Old and AD for (*left*) loops with CTCF loss at either both anchors, or (*center*) loops with CTCF loss at one anchor or (*right*) *in silico* generated CTCF loops as control. Notably, loops with CTCF loss also have loss in 3D interactions. **(e)** Scatterplot showing percentage of DEGs (Differentially Expressed Genes) vs CTCF fold-change for CTCF losses at loops. DEGs (q < 0.05) are represented separately for those exhibiting upregulation (shown in red) or downregulation (shown in blue) in AD vs Old. **(f-g)** GO analysis for genes at loops (< 5kb from anchor) with CTCF losses at either anchors showing **(f)** Biological Process and **(g)** Molecular Functions. **(h-i)** DNA motif enrichment analysis using HOMER at CTCF anchor sites with **(h)** stable CTCF or **(i)** CTCF loss. Top 5 motifs by significance are reported for panels h and i.

To investigate the correlation between CTCF peak losses in AD and changes in 3D chromatin interactions, we started by classifying loops based on the patterns of CTCF chIP-seq peak changes. This analysis yielded in four main types of loops (**Fig. 3B**): those with CTCF loss at both anchors (n=108); those with CTCF loss at only one anchor (n=2,790); those with CTCF gained at one anchor and stable at the other (n=955); and those with stable CTCF at both anchors (n=20,351), which, as expected, constituted the highest number of loops.

Subsequently, upon confirming a positive linear correlation between CTCF loop intensity and corresponding HiC intensity (**Fig. 3C**), we assessed the HiC signal intensity at sites with CTCF loop losses (a loss at either one anchor or both anchors) (**Fig. 3D**). This analysis showed that chromatin interactions, as measured by HiC signal intensity, were reduced at loop sites with loss in CTCF binding. This supports the association between CTCF losses and reductions in 3D chromatin interactions. Notably, when comparing the HiC signal at sites with random CTCF loops, no reduction in HiC signal was observed in AD, indicating that the decrease in chromatin interactions is specific to loops with CTCF losses.

Building upon our previous investigation into whether CTCF enrichment correlated with gene expression (**Fig. S2**), we conducted a similar analysis, this time focusing on the target genes in the 3D space (≤ 5 Kb from the other anchor) (**Fig. S4B**) rather than the adjacent gene in the linear space. Despite considering all genes expressed in AD or differentially expressed genes (DEGs) in AD versus Old, this analysis still uncovered no correlation between changes in CTCF and the magnitude of gene expression alterations (**Fig. S4B**). These findings further support the notion that CTCF levels are unlikely to serve as predictors of gene expression.

However, when considering CTCF peak losses at these loops and the percentage of associated DEGs (q < 0.05) (considering those ≤ 5Kb from either anchors), we found that the higher the loss of CTCF in AD, the lower the percentage of DEGs with upregulated genes, indicating that CTCF may be required for increased gene expression. We also found that the higher the loss of CTCF in AD, the higher the percentage of DEGs with decreased expression, reinforcing the view that CTCF is needed for maintaining the expression of these genes. These results are in agreement with findings correlating loop losses with DEGs in the CK-p25 mouse model of AD ^24^.

Overall, these findings underscore the key role of CTCF in mediating 3D chromatin organization and highlight its potential implications in AD pathology.

### Functional analysis of genes and regulatory elements linked to loops exhibiting CTCF loss: association with DNA methylation, RNA, and DNA damage genes

To gain further insight into the functional relevance of 3D genome conformation in AD, we investigated the genes and potential transcription factors associated with loops exhibiting CTCF losses in AD. Our analysis focused on genes located within 5 Kb from CTCF anchors on either side of a loop. For these genes, we analyzed those linked to a distal CTCF anchor with CTCF loss, thereby establishing a connection between distal CTCF and the target gene. Gene Ontology analysis performed in R revealed top Biological Process terms related to DNA methylation, RNA regulation and DNA damage checkpoint (**Fig. 3F**), as well as top Molecular Function terms related to ATP-dependent protein binding, snRNAs and transcription repression activity (**Fig. 3G**).

Next, we explored potential transcription factor binding at CTCF sites involved in these loops, distinguishing between anchor sites with CTCF losses and those with stable CTCF. Interestingly, while histone acetylation was stable at both anchors (**Fig. S4C-D**), in agreement with the analyses of CTCF peak losses (**Fig. S1C**), CTCF showed trends of reduced enrichment at sites considered as stable CTCF from the differential peak enrichment analyses. This finding suggests that significant loss of CTCF at one anchor may impact CTCF stability at the other anchor (**Fig. S4C-D**). DNA motif enrichment analysis using HOMER motif analysis confirmed CTCF binding sites and identified NEUROD1 and HIC1 as shared *Known* motifs for anchors with stable or loss CTCF (**Fig. 3H-I**).

BCL11A was specific to sites with stable CTCF, while ELK4 was specific to sites with CTCF loss. BORIS, the CTCF paralog expressed in testis, was expected to overlap with CTCF binding. *De Novo* motif analysis confirmed several of the findings from the *Known* motif analysis and highlighted new motifs such as PB0117.1, ZFP57, ETV2, among others (**Fig. 3H-I**). Overall, these results shed light on genes and potential transcription factors linked to CTCF loss in AD, indicating their potential involvement in loop dysregulation associated with the disease.

### CTCF loops define silencing domains and transcriptional repressors associated with AD

Given the remarkable changes in CTCF peaks in AD, we extended the studies to examine histone modifications associated with gene silencing, reasoning that the AD genome may be impacted by silencing marks together with structural changes. We performed H3K9me3 and H3K27me3 ChIP-seq in Young, Old and AD. We identified both gains and losses in H3K9me3 domains in AD vs Old, and the changes were predominantly gains in AD (data not shown). Interestingly, the gained H3K9me3 domains colocalized with CTCF chIP-seq changes were associated with downregulation of gene expression. The overall analysis of H3K27me3 did not show striking changes between AD and Old (data not shown).

We investigated potential regulatory factors including transcription factors that may be involved in the altered H3K9me3 domains and thus mediate the silencing of the associated genes. We integrated several analyses: the silenced genes that were associated with the H3K9me3 domains, CTCF HiChIP interactions, and histone acetylation changes. We performed DNA motif enrichment analysis at the upstream CTCF loop end, where enhancers are likely to be located, focusing on histone acetylation (H3K27ac/H3K9ac/H3K122ac combined) ChIP-seq peaks within the loop end. We found enrichment of several DNA binding motifs, of which a significant one was the DNA binding sequence of Pou6F1. Pou6F1 is a reported transcriptional repressor, and its expression is reported to increase in AD ^25^.

### Decrease of fly counterparts of CTCF and Pou6F1 promote Aβ42 neurodegeneration in a *Drosophila* model of AD

To extend our work to define the functional impact of some of these interactions, we first assessed if there were an effect to reduce the fly counterpart of CTCF in an Alzheimer’s fly model, expressing Aβ42. Reduction of CTCF using RNAi in the fly eye had no effect. However, co-expressing Aβ42 with reduced CTCF enhanced degeneration (**Fig 4**). These data support the idea that CTCF losses in AD are deleterious to the brain.

**Figure 4:**
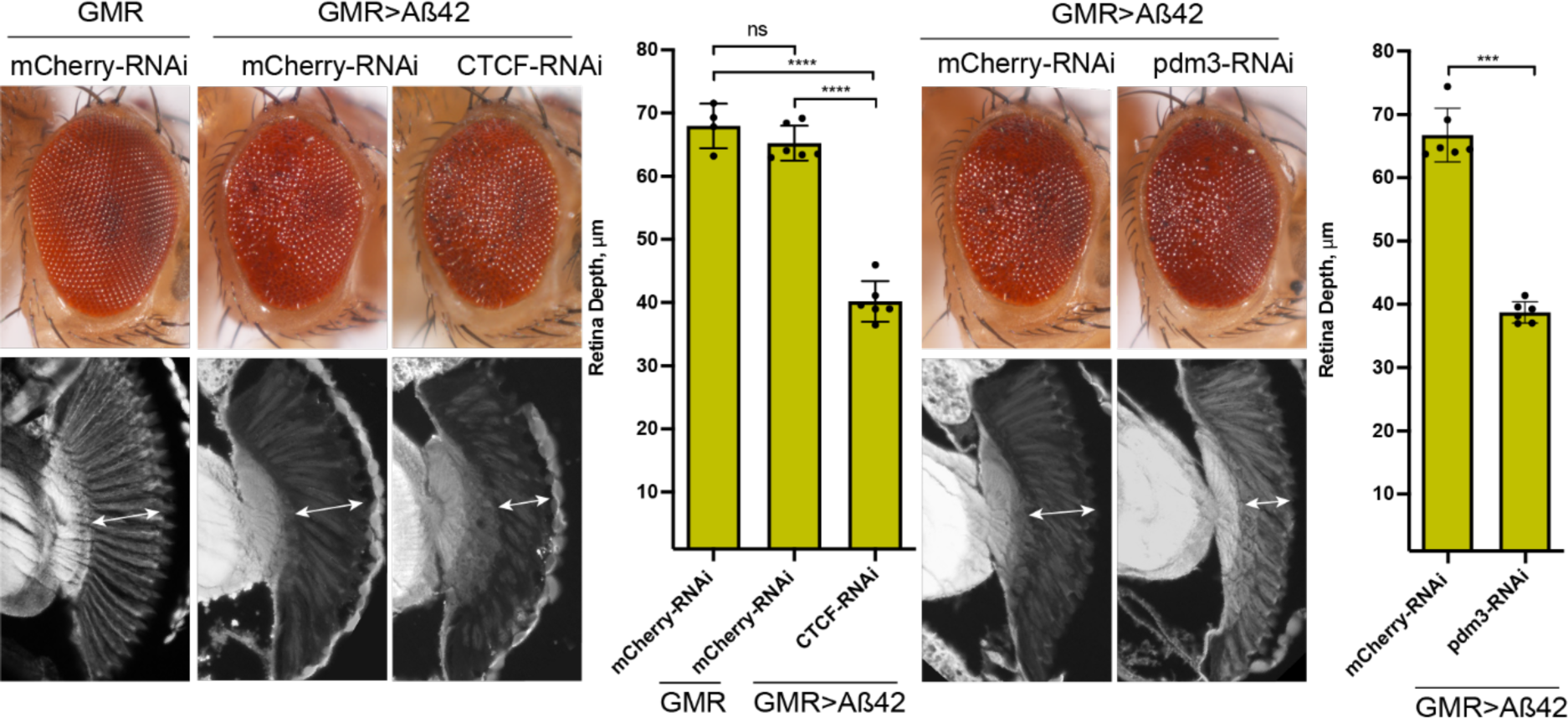
Reduction in CTCF and pdm-3 enhance Aβ42 toxicity in *Drosophila*. Expression of transgenes in the fly eye. Far left, normal eye. Expression of Aβ42 in the fly eye causes external disruption of the normally smooth eye, with no significant impact on the retinal depth (bottom). However reduction of CTCF leads to thinning of the retina; CTCF on its own has no effect (not shown). Similarly, reduction of pdm-3, a fly counterpart of Pou6F1, enhances toxicity by thinning the Aβ42 retina. This indicates that pdm-3 is normally protective. Pdm-3 loss on its own has no effect (not shown).

To define functional impact of Pou6F1, we investigated the impact of the fly ortholog of Pou6F1 again utilizing animals expressing amyloid-beta (Aβ42). The fly gene, *pdm3*, has been shown to function as a transcriptional repressor in the brain in *Drosophila* ^26^. Reduced expression via RNAi of *pdm3* led to enhancement of Aβ42-mediated toxicity (**Fig. 4**). These functional findings suggest that that the changes we detect in the AD genome with CTCF are deleterious, but the changes we detect associated with genome silencing that implicate Pou6F1, may be protective or homeostatic.

### AD eQTLs are enriched in CTCF losses in AD

Single nucleotide polymorphisms (SNPs) associated with disease tend to be enriched in non-coding regions, including regulatory elements such as enhancers, as well as potentially sites enriched for CTCF and RAD21. Given our findings indicating loss of CTCF and associated chromatin interactions in AD, we examined whether these genomic sites exhibit enrichment for AD GWAS SNPs or AD eQTLs, aiming to investigate a potential link between genetic risk for AD and epigenetic alterations associated with the disease. We also explored enrichment at RAD21 changing sites, although fewer RAD21 changes were observed in AD compared to CTCF (**see Fig. 1**). For consistency, we considered sites with peak changes (q< 0.05) in healthy aging (Old vs Young) in addition to those in AD (AD vs Old).

For the AD SNP enrichment analysis, we used an IGAP meta-analysis study ^27,34^. We employed a curated list of Alzheimer’s disease (AD)-associated SNPs (PL<L1L×L10^−5^) sourced from the International Genomics of Alzheimer’s Project (IGAP) meta-analysis study, which underwent two stages of clinical validation and included over 74,000 individuals. SNPs in linkage disequilibrium were merged into one region of analysis using PLINK v.1.9. For the AD eQTL enrichment analysis, we used a highly powered dataset with approximately 400 individuals containing eQTLs from the temporal cortex of AD cases (AD) (eQTLs, *n*L=L85,359), eQTLs from non-AD cases (other types of dementia; CTL) (eQTLs, *n*L=L156,134) and the two combined (BOTH; eQTLs, *n*L=L156,134) (as in ^9^). We employed INRICH ^28^ to assess the overlap between AD SNP or eQTLs and sites with peak changes. Focusing on the analysis of CTCF peak changes, INRICH analysis revealed no significant overlap between the AD SNPs and the CTCF changes in both healthy aging and AD (**Fig. 5A**). However, there was a significant overlap between CTCF losses in AD and eQTLs for all three classes of eQTLs (AD, *P*L=L9.79L×L10^−3^; CTL, P= 3.23L×L10^−2^; BOTH, *P*L=L1.99L×L10^−3^) (**Fig. 5A** and representations in **Fig. 5C-D**). In contrast, for RAD21, we identified no significance in any of the peak-SNPs enrichment analyses considering either healthy aging or AD changes for neither the AD SNP nor eQTLs (**Fig. 5B**).

**Fig. 5:**
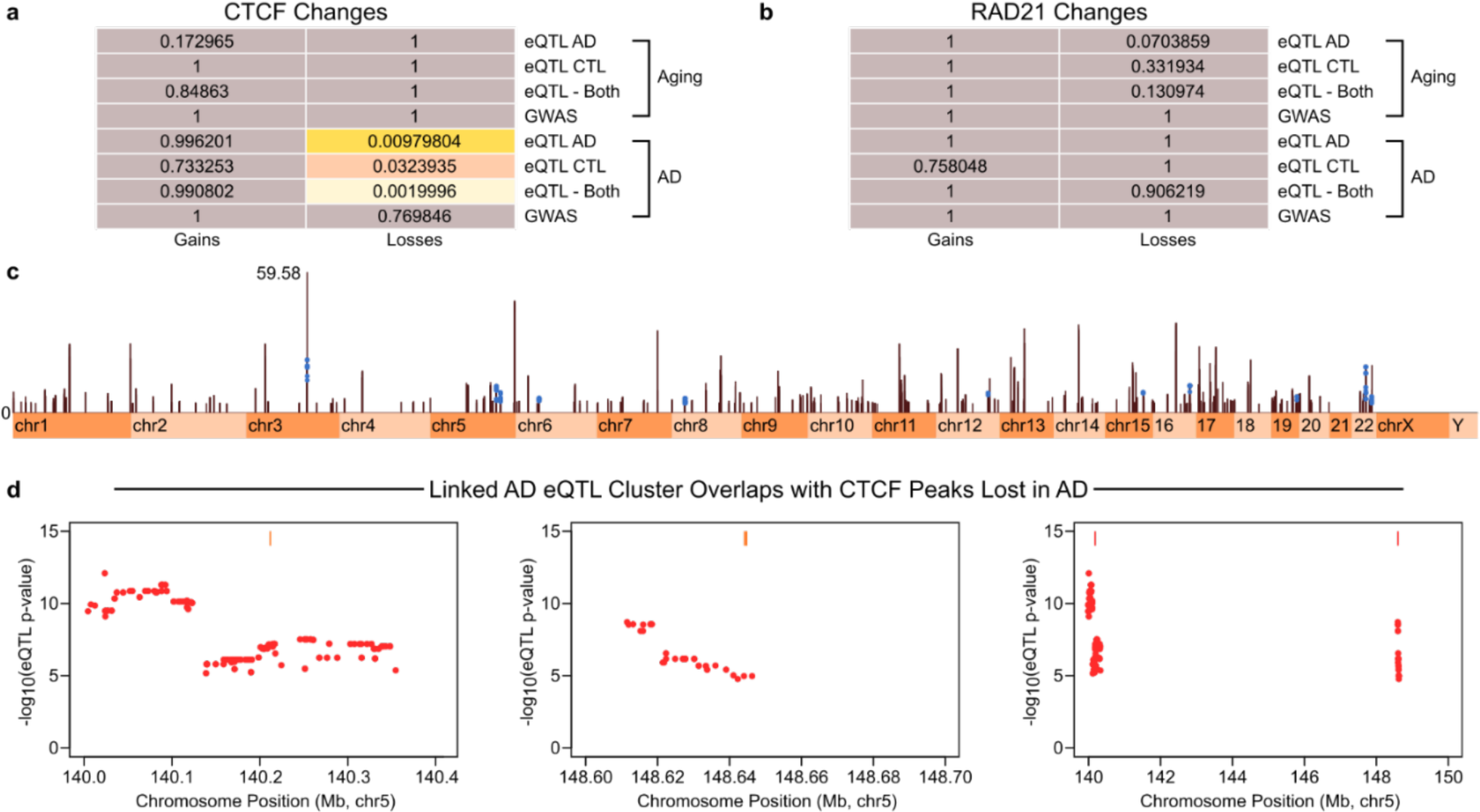
CTCF losses in AD are enriched in AD eQTLs. **(a)** Table reporting the significance of the overlap (p-value) between CTCF peak changes in aging or AD and AD GWAS SNPs or eQTLs linked to AD. **(b)** Table reporting the significance of the overlap (p-value) between RAD21 peak changes in aging or AD and AD GWAS SNPs or eQTLs linked to AD. INRICH was used for the enrichment analysis in (a) and (b). **(c)** Manhattan plot highlighting CTCF peak losses (blue dots) overlapping with AD eQTLs. **(d)** Representations of AD eQTLs overlapping with CTCF peak losses in AD (rightmost: zoomed-out view of two clusters, which are presented in detail in leftmost and middle panels).

Overall, these analyses further validate the significance of CTCF losses in AD and suggest a potential functional connection between genetic risk and CTCF loss-mediated alterations in AD.

## DISCUSSION

Here we have performed analysis of the genome of normal aging, by analyzing young and old brain tissue, and compared this with Alzheimer’s disease. Our study uses chromatin material isolated from the lateral temporal lobe, a region affected early in Alzheimer’s disease and associated with the memory loss. By including normal aging, and comparing this to AD, we have made some unexpected findings. First, our data in general indicate that CTCF remains in general largely stable in the genome with age (old compared to young), and by contrast the Alzheimer’s brain shows dramatically perturbed CTCF peaks, dominated by losses. Thus, whereas CTCF is in general stable with age, it is drastically affected by degeneration of Alzheimer’s disease. By contrast, RAD21 overall trends showed that, by comparison to CTCF, it remains largely stable with age and with disease. This could be because RAD21 has other functions in the genome, for example DNA repair ^17^. But it underscores that not all features of the genome are dramatically altered in Alzheimer’s vs normal. Moreover, we found that TADs were in general stable, with age and with disease. This is expected: TADs are thought to be highly stable structures ^7,19,20^. In investigating other changes and integrating the data, most of the changes we see are associated with the dramatic CTCF losses that occur in AD. *Drosophila* studies also indicate that reduction of CTCF leads to enhanced toxicity of amyloid beta protein, indicating that the dramatic impact on CTCF in AD is very likely deleterious.

Our analysis to date has been performed on bulk tissue, whereas single cell data, although of less read depth, will be highly informative. Such snRNA-seq data can address whether changes detected are characteristic of all cells, or are limited to neurons, or specific classes of neurons, or even seen in glia, which are emerging as critical players in Alzheimer’s disease ^29,30^. Our data and other’s data together ^8–10,12,13^ note that there are clearly robust epigenetic changes associated with the Alzheimer’s brain. Because we have included normal aging (old vs young), we can interpret some of the changes as being protective to the genome, as they are seen in healthy aging. Others, however, appear to be contributing to deterioration of the brain cells due to the disease, causing instability in the genome.

Key to all of these studies are questions related to how to protect the brain, how to enhance those changes that are characteristic of healthy aging of genome, and how to limit those that are associated with, or enhancing, deterioration of disease. Small molecules that impact epigenetic features of the genome are in development for cancer. Perhaps such approaches could be developed to enhance stability and healthy aging of the brain genome, thus to protect from disease.

## METHODS

### Brain tissue samples

Postmortem human brain samples from lateral temporal lobe (Brodmann area 21 or 20) were as previous ^8,9^ and obtained from the Center for Neurodegenerative Disease Research (CNDR) brain bank at the University of Pennsylvania (Penn). Informed consent for autopsy was obtained for all patients and it was approved by the Penn Institutional Review Board (Penn IRB). CNDR autopsy brain bank protocols are exempted from full human research (research on tissue derived from an autopsy is not considered human research – see https://humansubjects.nih.gov/human-specimens-cell-lines-data).

### ChIP-seq

ChIP-seq was performed as previously described ^8,9^. Briefly, nuclei were isolated from 200 mg frozen brain tissue (lateral temporal lobe) by dounce homogenization in nuclei isolation buffer (50 mM Tris-HCl at pH 7.5, 25 mM KCl, 5 mM MgCl2, 0.25 M sucrose and freshly added protease inhibitors and sodium butyrate) followed by ultracentrifugation on a 1.8 M sucrose cushion. Nuclei pellets were resuspended and crosslinked in 1% formaldehyde for 10 min at RT, quenched with 125 mM glycine for 5 min and washed twice in cold PBS. 2 x 10^6^ nuclei were lysed in nuclei lysis buffer (10 mM Tris-HCl at pH 8.0, 100 mM NaCl, 1 mM EDTA, 0.5 mM EGTA, 0.1% Na-deoxycholate, 0.5% N-lauroylsarcosine and freshly added protease inhibitors and sodium butyrate and) and chromatin was sheared using a Covaris S220 sonicator to a final ∼250 bp. Equal aliquots of sonicated chromatin were used per immunoprecipitation (IP) reaction with antibody CTCF and RAD21. ChIP reactions were incubated overnight at 4°C followed by ChIP washes and DNA was eluted in elution buffer (1% SDS, 50 mM Tris-HCl pH 8, 10 mM EDTA) at 65°C. Eluted DNA from ChIP and Input samples was reverse crosslinked and purified using PCR columns (Qiagen). 5 ng of ChIP and Input DNA was used to generate ChIP-seq libraries using the NEBNext Ultra DNA library prep kit for Illumina (New England Biolabs, NEB).

Libraries were multiplexed using NEBNext Multiplex Oligos for Illumina (dual index primers) and sequenced (75 bp; single-end) on a NextSeq 500 platform (Illumina) in accordance with the manufacturer’s protocol.

For HiC experiments, we followed the protocol of Rao et al ^31^. For HiChIP experiments, we followed the protocol of Mumbach et al ^22^:

### ChIP-seq analysis

CTCF and Rad21 chIP-seq analysis was performed largely as described ^8,9^. Briefly, demultiplexed ChIP-seq tags (∼20 million reads/sample per mark) were aligned to the human reference genome (assembly NCBI37/hg19) using Bowtie v1.1.1 allowing up to two mismatches per sequencing tag (parameters -m 1 --best). Reads mapped to mitochondria or ENCODE blacklist regions were removed from the analysis. Peaks were detected for each mark and in each sample using MACS2 with treatment-matched Input tags as control, as well as in each study group (Younger, Old and AD) by pooling reads of samples belonging to the same group. Peaks detected in the three study groups were filtered for peaks that were called in at least one sample. Comparison of histone acetyl enrichment across the three study groups was done using The MTL method. Briefly, MTLs were generated for each histone mark by merging peaks with 1 bp overlap across Younger, Old and AD. ChIP-seq signal in the MTL (unit: RPKM) was quantified for each individual sample using the Bwtool package (“bwtool summary”). In order to reduce the confounding variables due to changes in neuronal fractions across the samples, the top 10% MTLs with highest Pearson correlation with sample neuron fraction were masked from the analysis (neuron fractions for these samples were measured in our previous study ^8^). In order to cross-compare the different histone acetyl marks, masked MTLs for each H3K27ac, H3K9ac and H3K122ac mark were merged into a union set of acetylation sites (called “multiMTL”), and ChIP-seq signal for each histone mark and each individual patient was evaluated for these multiMTL sites (signal is only accounted for over each histone mark’s own MTLs). ChIP-seq signal over multiMTL sites was log_2_ transformed for downstream analysis (“log_2_[RPKM+1]”). Statistical significance of differential enrichments was assessed by performing a Wilcoxon rank-sum test (two-sided) for pairwise comparisons between study groups or 1-way ANOVA (two-sided) for a comparison of all three study groups.

For H3K9me3, H3K27me3, and chIP input, a different analysis workflow was used due to the tendency of these PTMs to form larger, domain-like structures. Bowtie2 v2.3.4.1 was used (parameters bowtie2 - p 32 --local -X 1000) to align the data to the hg19 / GRCh37 assembly of the human genome. Aligned tags were filtered using samtools v1.1 (parameters view-q 5-f 2), sorted by tag name (samtools sort with-n) and PCR deduplicated using PICARD MarkDuplicates v2.21.3-SNAPSHOT (parameters REMOVE_DUPLICATES=True ASSUME_SORT_ORDER=queryname). Tracks / bigWigs were created for visualization by first re-sorting BAM files by chromosome position (samtools sort) and then indexing using samtools index; following this, the chIP tracks were input-subtracted using deepTools bamCompare v3.4.3 with parameters--operation subtract. Enriched domains were called using RECOGNICER v1.0 with an FDR cutoff of 0.1. Differential domains were evaluated by first scoring domains for H3K9me3-input (tag density per kilobase) in each patient brain / sample, filtering out domains with no overlap to gene bodies / small domains (size < 10kb), then performing t tests to contrast study groups and FDR correcting by Benjamini-Hochberg step-up (also FDR 0.1).

### HiC Pre-Processing

Human 76bp paired-end reads were aligned independently to hg19 genome using bowtie2 ^32^ (global parameters:–very-sensitive –L 30 –score-min L,-0.6,-0.2 –end-to-end–reorder; local parameters:–very-sensitive –L 20 –scoremin L,-0.6,-0.2 –end-to-end–reorder) through the HiC-Pro software ^33^. Unmapped reads, non-uniquely mapped reads and PCR duplicates were filtered, uniquely aligned reads were paired, and replicates were merged. Cis-contact matrices were assembled by binning paired reads into uniform 20 kb bins. After matrix assembly, we normalized across samples using library size factors conditioned on genomic distance and then rounded the normalized matrices to the nearest whole number. Poorly mapped regions were removed based on hg19 75-mer CRG Alignability track from ENCODE. The interactions of 20kb bins were set to NaN if the average mappability of a 50kb window centered on that bin was below 50%. In addition, a high outlier filter was also implemented to further prevent balancing artifacts. We removed pixels in the raw contact matrices that exhibited high fold changes relative to a neighborhood of adjacent pixels after balancing. The neighborhood around a given pixel was defined by a 5 x 5 square footprint, which was used to determine a local median based on the values nearby pixels would have if they were to be balanced without the high outlier filtering step. If the value of a given pixel was greater than 4 times the local median or greater than 4 if the local median was less than 1, then that pixel was determined to be an outlier. Rows containing less than 35 non-zero pixels within 1.5 Mb of the diagonal were completely removed from the HiC analysis. After filtering, the remaining 20kb cis-contact matrices were balanced using Knight-Ruiz matrix balancing on merged replicates for every condition for each chromosome individually. The final bias factors were retained for subsequent loop calling.

### HiC Loop Calling

Expected modelling strategy similar to that used by the HiCCUPS algorithm proposed by Rao et al. ^31^ was implemented, with some alterations. For expected modelling and all subsequent loop calling steps, we restricted our analysis to bin-bin pairs with interaction distances of <= 20 Mb. All computations related to expected modeling were performed on the Knight-Ruiz balanced contact matrices binned at 20kb resolution.

To estimate the local background domain interaction frequency at each locus, donut expected model approach with parameter p=2, w=8 was utilized. For each matrix entry, the expected values were calculated using both the full donut window and just the lower-left region of the donut and the max of the two was taken i.e. expected = max(donut, lower-left). However, for bin-bin pairs with interaction distances within 300kb, we observed that the normal donut filters over-estimated the expected background, reducing the sensitivity of loop calling near the diagonal of the contact matrix. Therefore, to accurately capture short range interactions, we modeled the on-diagonal (< 300kb) background expected using only the upper-triangle of the donut footprint.

In summary, our matrices of final expected values were from the upper triangular corrected expected values for bin-bin pairs with interaction distances within 300kb and from the maximum of the donut and lower left footprints corrected expected values for bin-bin pairs with interaction distances beyond 300kb. Expected contact matrices were then deconvoluted back to discrete counts using the bias factor generated during Knight-Ruiz balancing. Each bin-bin pair in the rounded normalized matrices was tested for significance under a Poisson distribution parameterized by its corresponding deconvoluted expected values. To perform multiple testing correction, we applied the lambda-chunking strategy proposed by Aiden and colleagues. Briefly, we first stratified bin-bin pairs according to their biased expected values using logarithmically spaced bins with a bin spacing 2^1/3^. We then applied Benjamini-Hochberg false discovery rate control for the p-values for each chunk separately to obtain q-value matrices. For an interaction to be called as significantly enriched above the background, it was required to pass 3 thresholds: 1) a q-value threshold < 0.1 (corresponding to an FDR of 10%); 2) a balanced contact value ≥ 8.0; 3) a fold-change between balanced contact value and final expected value ≥ 1.5.

Matrix entries composed of directly adjacent bin-bin pairs passing these three thresholds were clustered into loops. We applied a progressive stringent q-value thresholding and re-clustered bin-bin pairs that passed the more stringent q-value. The q-value thresholds were applied in order-of-magnitude steps from 0.1 to 1e-20 FDR. This helped in refining our initial clusters that sometimes included multiple visually-apparent loops into one large supercluster to now have smaller clusters that represent sub-regions of relatively higher significance. To further remove the possibility of false-positives, we removed clusters with fewer than three adjacent significant bin-bin pairs and whose interaction distance was within 60kb from the diagonal.

### Target Gene Analysis

We used the loop calls described in the previous section and linkage disequilibrium (LD) blocks of 32 single nucleotide variants (SNVs) identified as statistically associated with Alzheimer’s Disease ^34^ to identify target genes. The rationale for using LD blocks is that the causal SNV is likely not genotyped by Kunkle et al. We computed LD blocks from 1000 Genomes Project Phase 3 variant calls. First, we filtered the calls to only include CEU, FIN, GBR, IBS, and TSI European populations. Then, we left-aligned the variants and split multi-allelic variants into separate variants using BCFTOOLS NORM. Duplicate variants and CNVs were removed, and ambiguously named SNVs (such as “.”) were renamed according to their location using custom scripts. Next, we extracted LD blocks defined by an r^2^ threshold of 0.3 using PLINK v1.90b6.18 with the arguments--vcf--r2--ld-window-kb 1000--ld-window-r2 0.3--ld-window 99999--ld-snp-list.

For our “conventional approach” for identifying target genes, we directly intersected the SNVs in the LD blocks with a genome-wide list of hg19 RefSeq genes. This list was downloaded from the UCSC Genome Browser on March 24, 2021 and was modified to include 5kb promoter regions. For our “looping approach” for identifying target genes, each loop call was first parsed into separate anchors and then intersected with LD blocks. If a loop anchor intersected with one or more SNVs in an LD block, then the opposite anchor was intersected with the list of hg19 RefSeq genes.

### TAD and subTAD detection using 3DNetMod

TADs were identified genome-wide using 3DNetMod ^18,35,36^ on the Knight-Ruiz balanced and scaled Hi-C matrices at 20 kb resolution. For each condition, log-transformed genome-wide counts data were used and the following parameters in 3DNetMod settings file were set for TAD calling: diagonal_density=0.65, consecutive_diagonal_zero=62, pleateau=1, chaosfilter=True, chaos_pct=0.85, num_part=20, size_threshold=2, size_s1=4000000, size_s2=12000000, var_threshold=100, boundary_buffer=80000. If there was an overlap between adjacent domain boundaries such that boundaries were within +/-70kb of each other, then domains were merged into a single TAD/subTAD separated by the largest genomic distance start and end coordinates. Subsequently, domains were classified into three layers TADs, subTADs and ultra-nested subTADs as previously described^18,36,37^. A non-redundant genomewide list of TAD/subTAD boundaries were generated from the domain calls by separating their start and end coordinates for each of the 3 conditions.

### Differential boundary analysis

Likelihood ratio test was performed to determine genome-wide differential boundaries between AD vs. Old and Old vs. Young pairwise comparisons. Briefly, each of the boundaries were padded +/-1Mb on either side. From the Knight-Ruiz scaled, balanced 20kb binned Hi-C matrices for each replicate separately per condition, mean insulation score was computed for each boundary window. To compare Old vs. Young, the null hypothesis was that the mean boundary strength in Old and Young was the same.

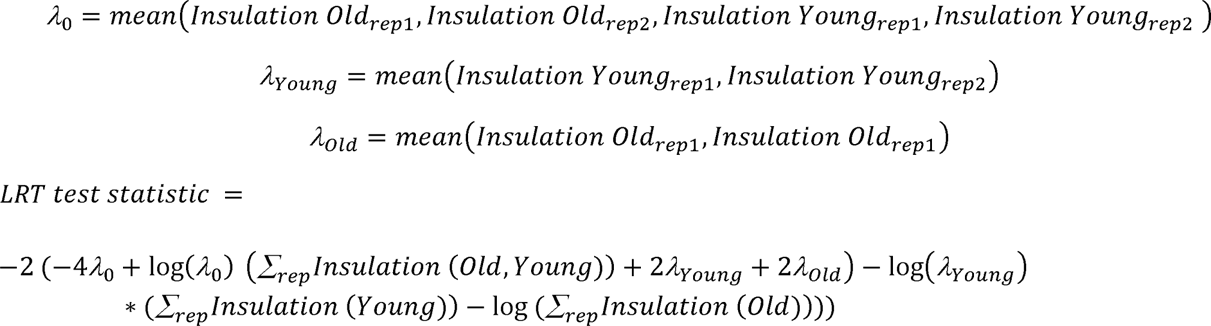

Finally, for each test statistic, *p-value* was computed using stats.chi2.sf(LRT_statistic, 1) scipy stats in python. We used p-value < 0.05 to be called a differential boundary. Thus for each boundary genome-wide, change in boundary strength was computed as 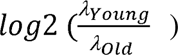 and average strength as 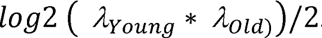.

Similarly, differential boundaries between AD vs. Old comparison were computed.

### Computing Insulation score (IS)

Insulation score genomewide for AD, Old and Young conditions were computed as described previously ^18,35,36^. On the Knight-Ruiz balanced, scaled 20kb binned Hi-C matrices, a 6 kb square window without any offset from the diagonal was applied. Within the 6 x 6 IS window counts were summed, normalized by mean of the chromosome-wide and finally counts were log transformed. Mean insulation score +/-1Mb each boundary was computed for each of the 3 conditions.

### RNA-seq analysis

RNA-seq data were previously generated ^8^ and processed for the same brain samples (lateral temporal lobe) analyzed in this study. Briefly, total RNA was isolated using the RNAeasy Mini kit (Qiagen) and treated with RNase-free DNase step (Qiagen). Ribosomal RNA was removed using the rRNA Depletion kit (NEB) and multiplexed RNA-seq libraries were generated using the NEBNext Ultra Directional RNA library Prep Kit for Illumina (NEB). Libraries were sequenced (75 bp, single-end) on a NextSeq 500 platform (Illumina) in accordance with the manufacturer’s protocol. RNA-seq reads were aligned to the human reference genome (assembly GRCh37.75/hg19) using STAR with default parameters.

FeatureCounts was used to generate a matrix of mapped fragments per RefSeq annotated gene. Analysis for differential gene expression between AD and Old was performed using DESeq2 R package with FDR < 0.05. For correlation between ChIP-seq and RNA-seq data, DESeq2 normalized read count over each gene was normalized by per Kb of gene length and then transformed to log2 space (“log2[Count+1]”, where “Count” is the average read count of each study group). Peaks were linked to the nearest annotated gene in the RefSeq database.

### Genome Browser tracks

Generation and visualization of ChIP-seq tracks was conducted as follows. For each individual sample and the three study groups (Young, Old and AD; pooled reads of samples from the same group), tag pileup profiles (bedgraph files) were generated during MACS2 peak calling process to visualize signal per million reads (RPM). Treatment-matched input signal was subtracted from the pileup profiles using MACS2 (“bdgcmp”). Finally, bigwig files were generated using UCSC toolkit (“bedGraphToBigWig”) and uploaded on the UCSC Genome Browser.

### Functional analyses

Functional analysis of genes differentially expressed between AD and Old was performed using DAVID ^37^. For GREAT analysis, the “single nearest gene” rule was used to associate peaks to RefSeq genes and Biological Process (BP) terms were filtered to be significant by both the binomial and hypergeometric tests (FDR < 5%) and binomial fold enrichment filter of 2. For DAVID analysis, each peak was linked to its nearest RefSeq gene and significant terms (FDR < 10%) for BP, Cellular Component (CC), Molecular Function (MF) and Tissue are reported. FDR < 10% represents the threshold of significance in DAVID.

### Graphical representation

Scatter plots and boxplots of ChIP-seq data were visualized using R package ggplot2 (version 3.3.1). Metaplots and signal heatmaps centered around peaks were generated using Deeptools (“computeMatrix” and “plotHeatmap”, version 2.5.7).

### Association between peaks and AD GWAS SNPs

INRICH was used to infer the relationship between peaks and PLINK-joined AD GWAS SNP intervals (linkage due to HapMap release 23) using standard parameters. The set of all changed peaks (*P* < 0.05, 1-way ANOVA), was the background for the experiment.

### eQTL data processing and sampling analysis

eQTL data tables were downloaded and used as previously described ^9^. For each dataset, custom scripts, also available by request, were used to summarize the overlap counts in easy to parse files that were then read into the R programming language which was used to perform the empirical enrichment analyses.

### Drosophila studies

Flies were maintained on standard yeast/cornmeal-based medium at 26°C on a 12hr:12hr light–dark cycle. *gmr*-Gal4 (#1104), UAS-*mCherry*-RNAi (#35785), UAS-*CTCF-*RNAi (#40850), UAS-*pdm3-*RNAi (#53887) transgenic fly lines were obtained from the Bloomington *Drosophila* Stock Center; the β-amyloid fly line was generously shared by P. Fernandez-Funez (University of Minnesota). *Drosophila* eye and retinal imaging were performed as described^8^. Images of retina sections were analyzed by ImageJ to measure the retinal depth across a consistent plane of the brain at the point of the optic chiasm. For each fly, 4 sections in the region of interest were measured and the average was used as the retinal depth for that animal. 4-6 animals/genotype were analyzed.

## Data & Code availability

The data that support the findings of this study will be available through the NCBI Gene Expression Omnibus (GEO) repository. Code developed for the analyses performed in this study will be made available.

---

**Fig. S1:**
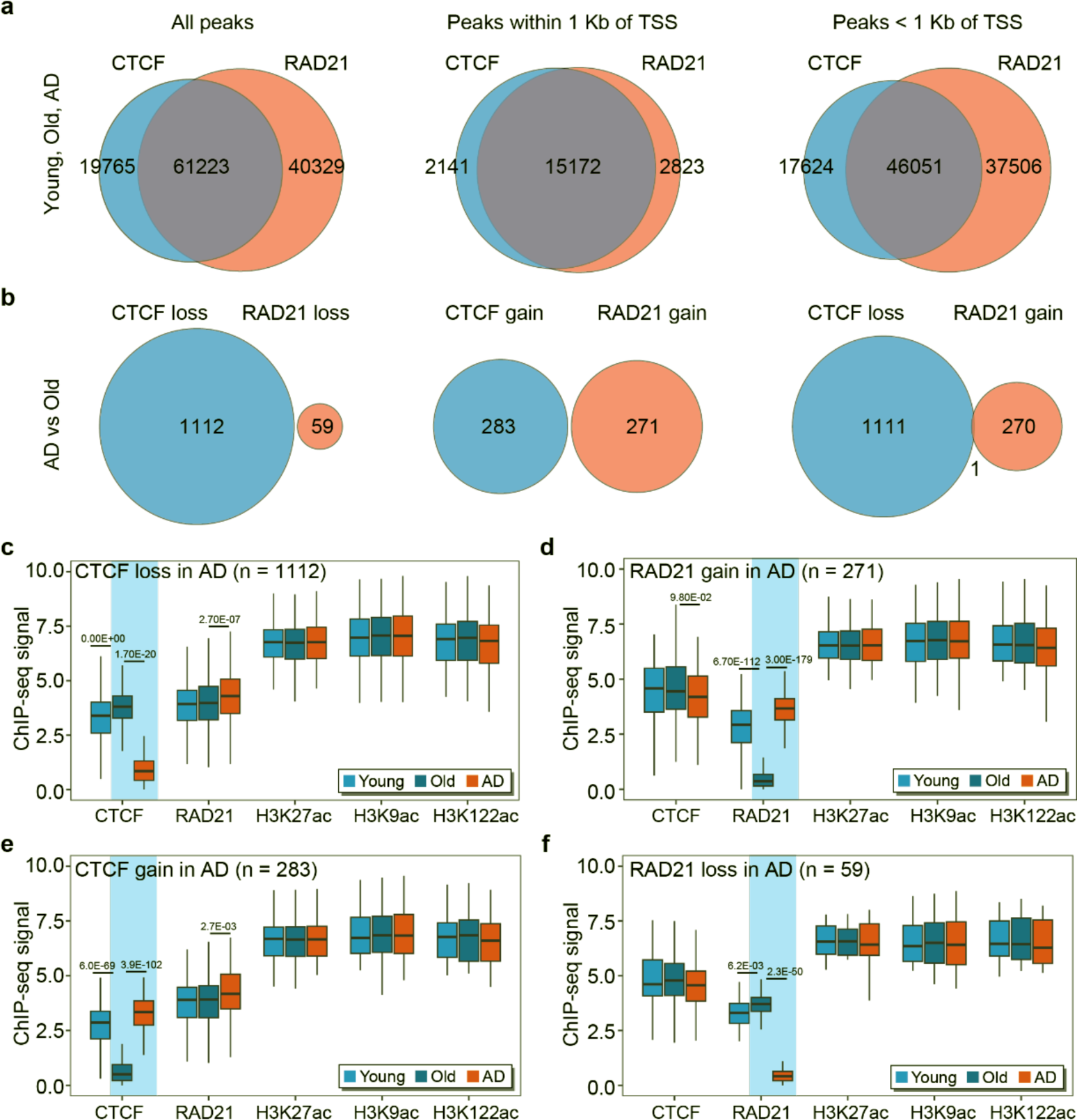
CTCF and RAD21 peak changes in AD and comparison with healthy aging. **(a)** Venn diagram showing CTCF and RAD21 peak overlap considering (*left*) all CTCF and RAD21 peaks, or (**center**) CTCF and RAD21 peaks at the promoter (≤ 1 Kb), or (*right*) CTCF and RAD21 peaks outside the promoter (> 1 Kb) such as enhancers for peaks detected in Young, Old or AD (union of peaks). **(b)** Venn diagram shown the overlap between CTCF and RAD21 peak changes for (*left*) peaks with both CTCF and RAD21 losses, (*center*) peaks with both CTCF and RAD21 gains, or (*right*) peaks with CTCF loss and RAD21 gain in AD vs Old. **(c-f)** Boxplot showing CTCF, RAD21, H3K27ac, H3K9ac and H3K122ac enrichment in Young, Old and AD for genomic sites with **(c)** CTCF loss, **(d)** RAD21 gain, **(e)** CTCF gain, or **(f)** RAD21 loss in AD vs Old. P-values are reported for statistically significant differences in ChIP-seq signal.

**Fig. S2:**
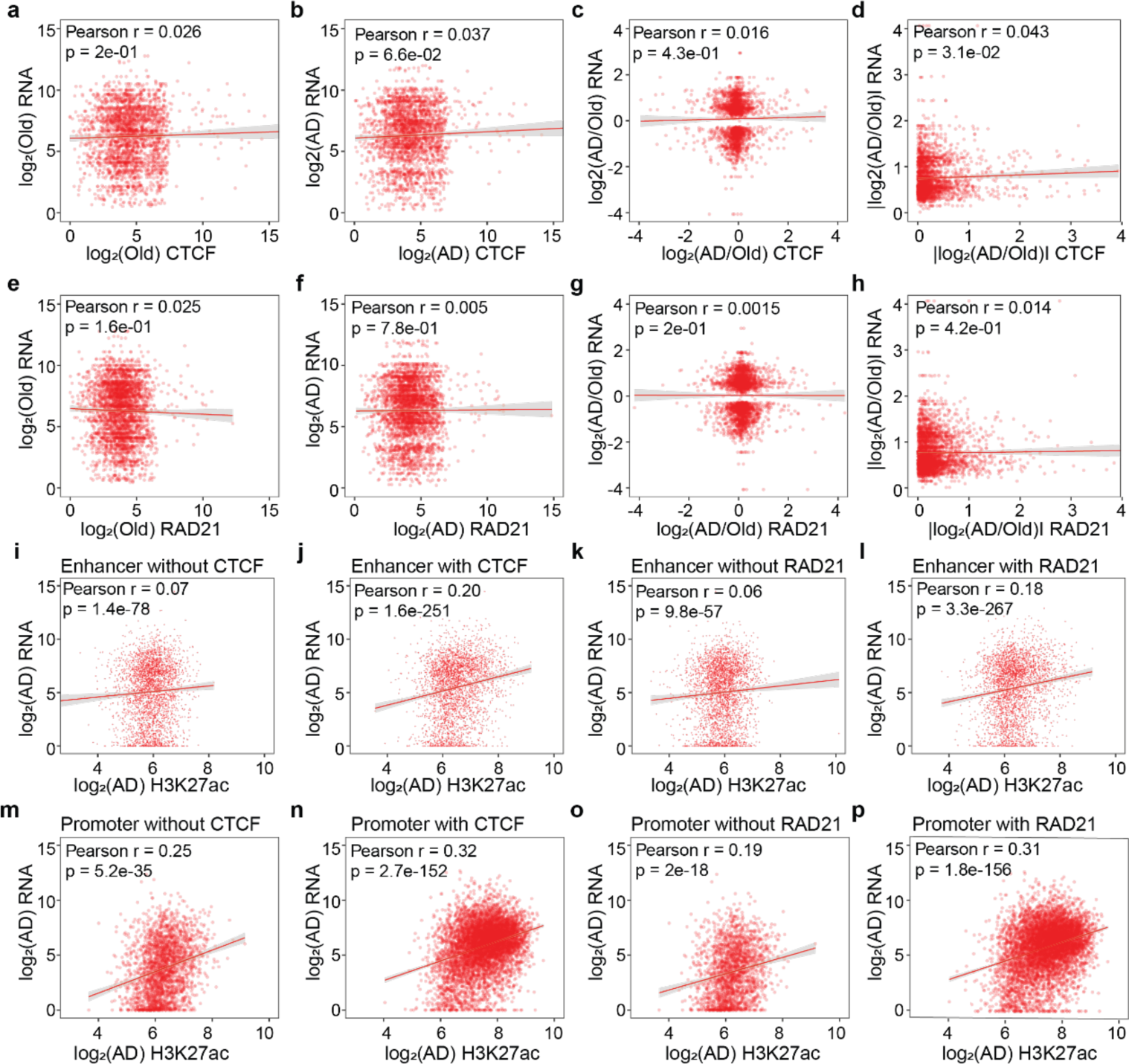
Genome-wide correlations between ChIP-seq and RNA-seq for regulatory elements enriched with CTCF or RAD21. **(a-b)** Scatterplot showing CTCF peak enrichment vs RNA-seq of nearest gene in **(a)** Old and **(b)** AD. **(c-d)** Scatterplot showing CTCF peak fold-change vs RNA-seq fold-change of nearest gene in AD vs Old considering **(c)** positive and negative fold-change or **(d)** the amplitude of change reported as absolute value. **(e-f)** Scatterplot showing RAD21 peak enrichment vs RNA-seq of nearest gene in **(e)** Old and **(f)** AD. **(g-h)** Scatterplot showing RAD21 peak fold-change vs RNA-seq fold-change of nearest gene in AD vs Old considering **(c)** positive and negative fold-change or **(d)** the amplitude of change reported as absolute value. RNA-seq is measured for DEGs (differentially expressed genes, q < 0.05) throughout panels a-h. **(i-l)** Scatterplot of H3K27ac enrichment vs RNA-seq of nearest gene in AD for enhancers (> 1Kb from TSS) **(i)** without or **(j)** with CTCF, or enhancers **(k)** without or **(i)** with RAD21. 3,000 randomly chosen points are shown in each panel. **(m-p)** Scatterplot of H3K27ac enrichment vs RNA-seq of nearest gene in AD for promoters (≤ 1Kb from TSS) **(i)** without or **(j)** with CTCF, or promoters **(k)** without or **(i)** with RAD21. Pearson’s correlation coefficient and p-value are indicated in the panel.

**Fig. S3:**
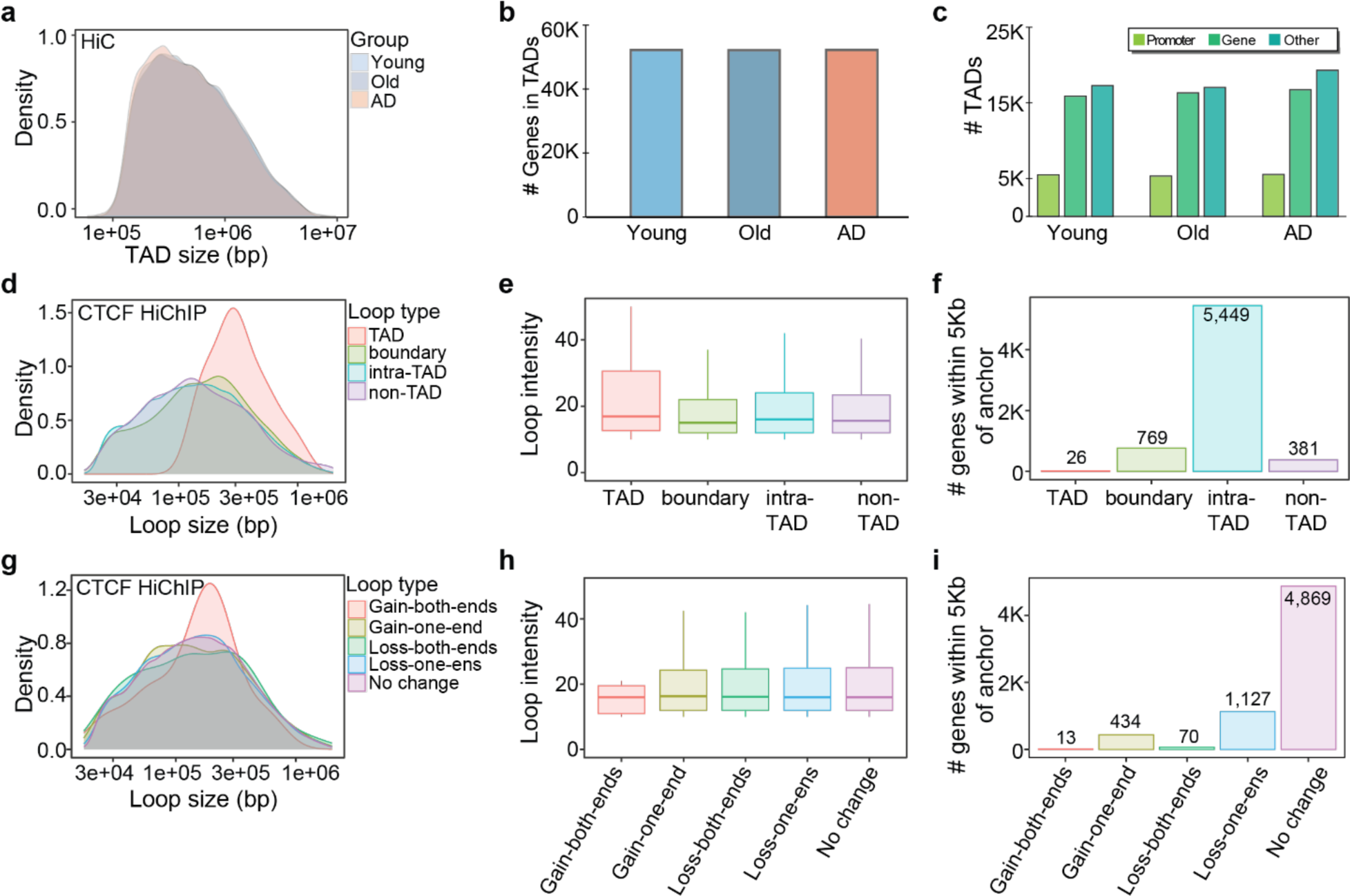
Genome-wide characterization of HiC TADs and CTCF HiChIP loops. **(a)** Density plot showing size distribution of TADs (bp) in Young, Old or AD. **(b)** Barplot showing number of genes contained in TADs in Young, Old or AD. **(c)** Number of TADs which contain promoters, genes or other genomic elements in Young, Old or AD. **(d-f)** Genome wide-analysis of CTCF HiChIP loops showing **(d)** density plot of loop size distribution, **(e)** boxplot of loop intensity and **(f)** number of genes within loops (≤ 5 Kb from anchor) for loops categorized based on their location relative to TADs (TAD, boundary, intra-TAD or non-TAD loops). **(g-i)** Genome wide-analysis of CTCF HiChIP loops showing **(g)** density plot of loop size distribution, **(h)** boxplot of loop intensity and **(i)** number of genes within loops (≤ 5 Kb from anchor) for loops categorized based on CTCF peak changes in AD vs Old at each anchor point (gain at both ends, gain at one end, loss at both ends, loss at one end, stable CTCF). Notably, loops categorized based on CTCF changes at anchors were considered for downstream functional analyses.

**Fig. S4:**
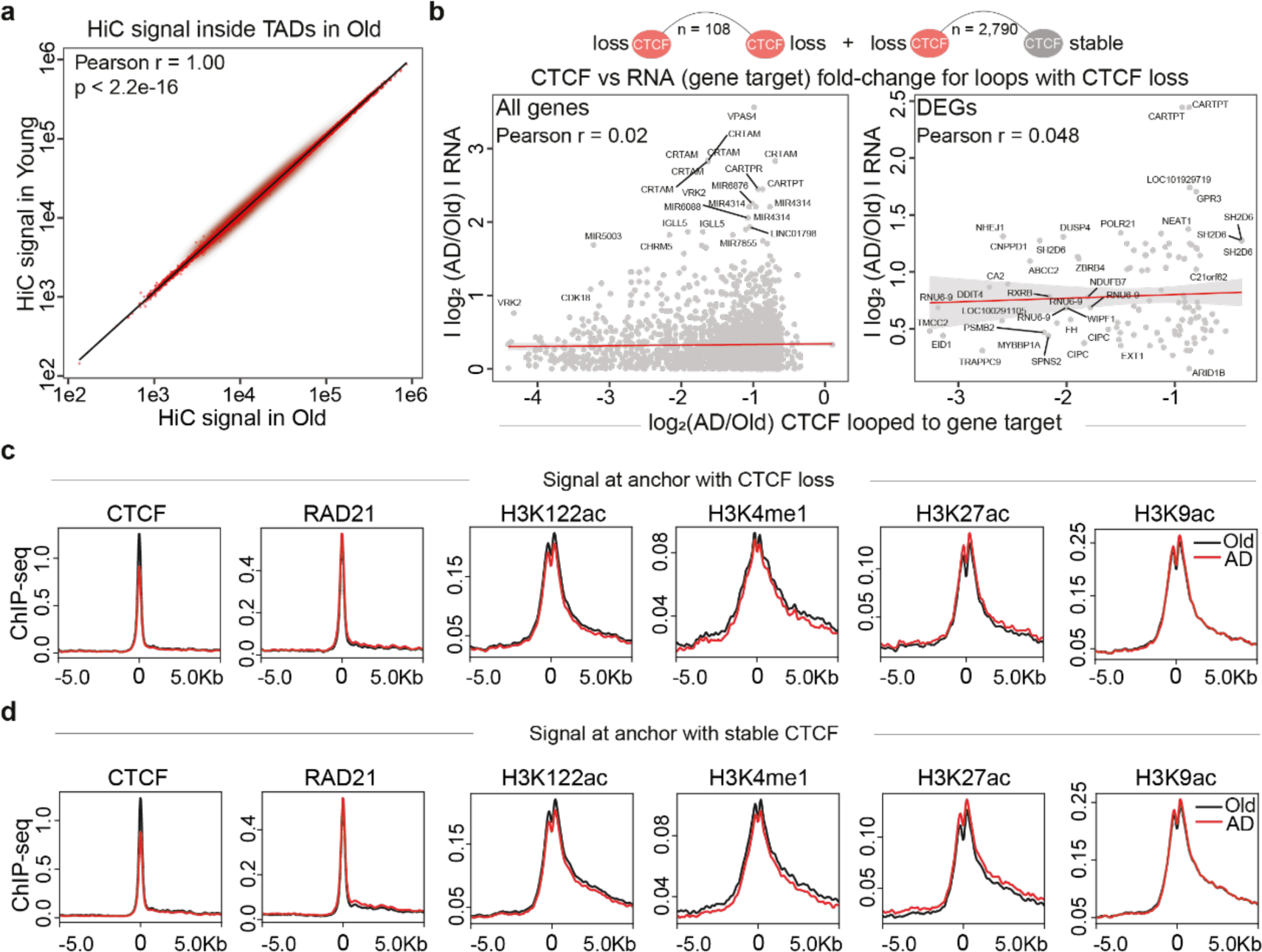
Extended analysis of HiC and CTCF HiChIP loops in healthy aging and AD. **(a)** Scatterplot showing HiC signal in Old vs Young for TADs detected in Old. Linear regression trendline, Pearson’s correlation coefficient and p-value are indicated in the panel. **(b)** Scatterplot showing CTCF peak fold-change vs absolute RNA-seq fold-change for CTCF loops with CTCF losses in AD vs Old considering CTCF anchors and their gene target. The analysis is based on considering (*left*) all genes or (*center*) DEGs (differentially expressed genes) with which CTCF interacts with. Linear regression trendlines, Pearson’s correlation coefficients and p-values are indicated in the panel. Illustration of CTCF loops with CTCF losses at either one or both anchor sites reported on top. **(c-d)** Metaplots showing CTCF, RAD21, H3K122ac, H3K4me1, H3K27ac and H3K9ac enrichment at loop anchors with **(c)** CTCF loss or **(d)** stable CTCF in AD vs Old. Metaplots are centered at the CTCF anchor site.

